# End-to-end mapping of membrane transport from chemical structure to microorganisms

**DOI:** 10.64898/2026.05.12.724480

**Authors:** Guillaume Gricourt, Thomas Duigou, Philippe Meyer, Jean-Loup Faulon

**Author notes:** Correspondence and requests for materials should be addressed to Jean-Loup Faulon.

## Abstract

Membrane transport is a fundamental biological process with profound implications for pharmacology, biotechnology, and microbiology. While computational approaches have largely adopted a protein-centric perspective to annotate transportomes, inferring transport function directly from the intrinsic properties of substrates remains a major challenge. Addressing transport at the compound level enables the systematic evaluation of whether molecules undergo active transport and by which mechanisms, independent of prior transporter annotation. Here, we introduce ChemProFlow, a comprehensive computational framework that redefines transport analysis from a substrate-centric perspective. By integrating geometric deep learning with orthology-based genomic mapping, ChemProFlow predicts molecular transportability, assigns transport mechanisms according to the Transporter Classification Database, and identifies the microorganisms encoding the corresponding transport systems. We show that this integrated pipeline enables scalable, end-to-end mapping of substrate-transporter-organism relationships, with broad applications in pharmacology for anticipating drug transport, in biotechnology for guiding strain engineering, and in microbiology for dissecting substrate utilization across diverse taxa. By capturing the chemical derminants of transportabiliy, ChemProFlow generalizes to previously unseen substrates and provides a high-throughput framework for systematic exploration of molecular transport across diverse biological contexts.

## Introduction

Membrane transport is a fundamental biological process that enables the selective translocation of molecules across lipid bilayers. Transfer between the external compartment and the cell occurs either passively, following a concentration gradient, or actively via channels and carrier proteins^1^. In clinical pharmacology, these transporters are key determinants of drug disposition and efficacy, systemic toxicity, and complex Drug-Drug interactions^2^. Beyond pharmacology, membrane transport has emerged as a central focus in biotechnology. Transport proteins are increasingly leveraged to enhance the uptake of pollutants in bioremediation applications^3^. In parallel, characterizing exchanged metabolites is critical for the rational design of microbial communities^4^, a rapidly expanding field with broad applications in environmental, industrial, and biomedical contexts^5^. Furthermore, transporter engineering has arisen as a key strategy for increasing microbial production yields in engineered cell factories^6^. Collectively, these developments highlight the growing need for scalable and systematic approaches to identify substrate-transporter relationships across diverse biological and technological contexts.

Transport proteins, together with their associated transport reactions, are manually curated and catalogued in the Transport Classification database (TCDB)^7^. Each transporter is assigned a Transport Classification Identifier (TC-ID), a five-level hierarchical code that organizes transporters by their taxonomic group, protein family relationships, and substrate specificity, similar in concept to the Enzyme Commission (EC) numbering system^8^. EC numbers, which provide functional insight through a hierarchical classification of catalytic activity, have been extensively used in biocatalysis to infer enzyme function and predict biochemical reactions, especially when integrated with artificial intelligence algorithms^9^. Similarly, TC-IDs offer a hierarchical framework for membrane transport proteins that supports inference of transport mechanisms^10^.

To infer transporter function, numerous studies have aimed to distinguish transporter proteins from non-transporters using sequence-based computational methods, adopting a protein-centric perspective. For example, TransATH^11^ improved upon the GBlast^12^ algorithm by integrating heuristic rules to develop an end-to-end transporter annotation pipeline, combining the TCDB database with homology search principles based on Blastp. This homology-based approach was also implemented in gapseq^13^, a tool designed to reconstruct Genome Scaled Metabolic Models (GEMs) from genomes. More broadly, homology-driven strategies constitute a central component of similarity-based ortholog detection methods, which aim to infer functional conservation between proteins across species^14^. These approaches are computationally efficient and readily scalable, frequently relying on the Reciprocal Best Hit (RBH) heuristic^15^. This heuristic identifies candidate orthologs by selecting sequence pairs of sequences from two distinct genomes that represent the top-ranking alignment match for one another, assuming that mutual sequence similarity reflects a shared evolutionary origin^16^. Beyond homology-driven strategies, TooT-T^17^ and, more recently, DEEPTRANS^18^, have improved transporter protein identification by leveraging advanced machine learning and transformer-based architectures.

Machine learning has also been applied to substrate class prediction directly from protein sequences. TranCEP^19^ employed sequence-derived features combined with support vector machines to classify substrates such as sugars and ions, whereas TooT-SS^20^ leveraged transformer-based representations to associate transporter proteins with specific substrate categories. Denger and Helms^21^ further explored the classification of substrates using gene ontology in a multi-labeled learning framework. More recently, SPOT^22^, integrated into the DeepMolecules^23^ webserver, demonstrated impressive performance in predicting protein sequence-substrate associations using advanced deep learning techniques. To construct their dataset, the authors paired each substrate with its corresponding transporter to form positive data points and generated negated examples by randomly associating substrates with unrelated proteins, thereby enabling the model to distinguish between related and unrelated pairs. In contrast, Drug-Drug interactions studies avoid random negative sampling, which assumes that all unobserved associations are negative, an assumption that may not hold for biological data^24^, and instead adopt a Positive-Unlabeled (PU) learning strategy that does not require explicit negative examples^25^.

While substantial progress has been made in protein-centric approaches to transporter prediction, relatively few studies have attempted to infer transport function directly from the intrinsic properties of substrates themselves. Early efforts relied on molecular descriptors combined with traditional machine-learning models to identify fragments associated with a limited number of transporters of interest^26^. More recently, the MONSTROUS web server leveraged Geometric Deep Learning (GDL) to predict interactions between small molecules and a restricted set of transporter proteins^27^. For instance, the widely adopted ChemProp, a GDL framework, has demonstrated strong performance across a broad range of molecular modelling tasks^28,29^. However, these approaches focus on predefined subsets of transporters and do not address the complementary and largely unexplored question central to a substrate-centric perspective: whether a given molecule is transportable and by which mechanisms.

Despite the rapid development of methods for annotating protein transporters, a fundamental asymmetry remains: transport function is typically inferred from proteins to substrates rather than from substrates to proteins. This limitation constrains our ability to evaluate the transport potential of novel or poorly characterized molecules, particularly when corresponding transporters are unknown or unannotated. A substrate-centric framework would instead begin from molecular structure, infer transportability and mechanism, and subsequently connect these predictions to genomic context. Such an approach would complement sequence-based strategies and extend transport analysis beyond the constraints imposed by existing protein annotations.

To address this gap, ChemProFlow is designed to: (i) determine whether a molecule is involved in a transport mechanism, (ii) infer its transport function by predicting the corresponding TC-ID and, (iii) identify the transportome of microorganisms through homology-based searches using the RBH method. When integrated, these modules enable the retrieval of microorganisms associated with specific transported substrates and the characterization of the underlying transport mechanisms. The framework provides an end-to-end solution capable of processing large datasets, enabling high-throughput exploration of molecular transport properties and assessment of transport potential prior to protein-level characterization.

## Results

### The ChemProFlow architecture

The ChemProFlow framework integrates several interconnected modules designed to predict and interpret molecular transport processes, beginning with a molecule’s structure and culminating in the identification of associated transport proteins and potential host organisms (Figure 1). The first module constructs substrate datasets for model training by integrating information from Rhea, the TCDB, and PubChem (see “Transport substrate datasets” section of Methods). The second module takes a molecular substrate, either drawn from these datasets or provided by the user, and generates a molecular graph embedding, which captures the topological and chemical features of the molecule (refer to “Molecular embedding” section of Methods). This embedding is then processed in a third module by a Graph Neural Network (GNN) trained using a PU learning strategy, followed by a post-hoc calibration step to enhance the reliability of predicted probabilities. The fourth module extends this analysis by employing a multi-label GNN model to predict the TC-ID for the query molecule, acknowledging that a single substrate can participate in multiple transport mechanisms. Lastly, the fifth module integrates these molecular predictions with genomic context to identify microorganisms, currently focusing on bacteria and fungi, that harbour transport mechanisms for the query molecule, based on the RBH principle. By cross-referencing the TC-IDs predicted in the third module, this component identifies specific taxa possessing cellular transport processes dedicated to the initial substrate. Together, these modules constitute an end-to-end substrate-centric workflow that bridges molecular features, transport classification, and genomic evidence within a unified predictive framework. In summary, the complete pipeline determines the transportability of a substrate, assigns it to a defined cellular mechanism, and retrieves the organisms likely to encode the relevant transport proteins.

**Figure 1.**
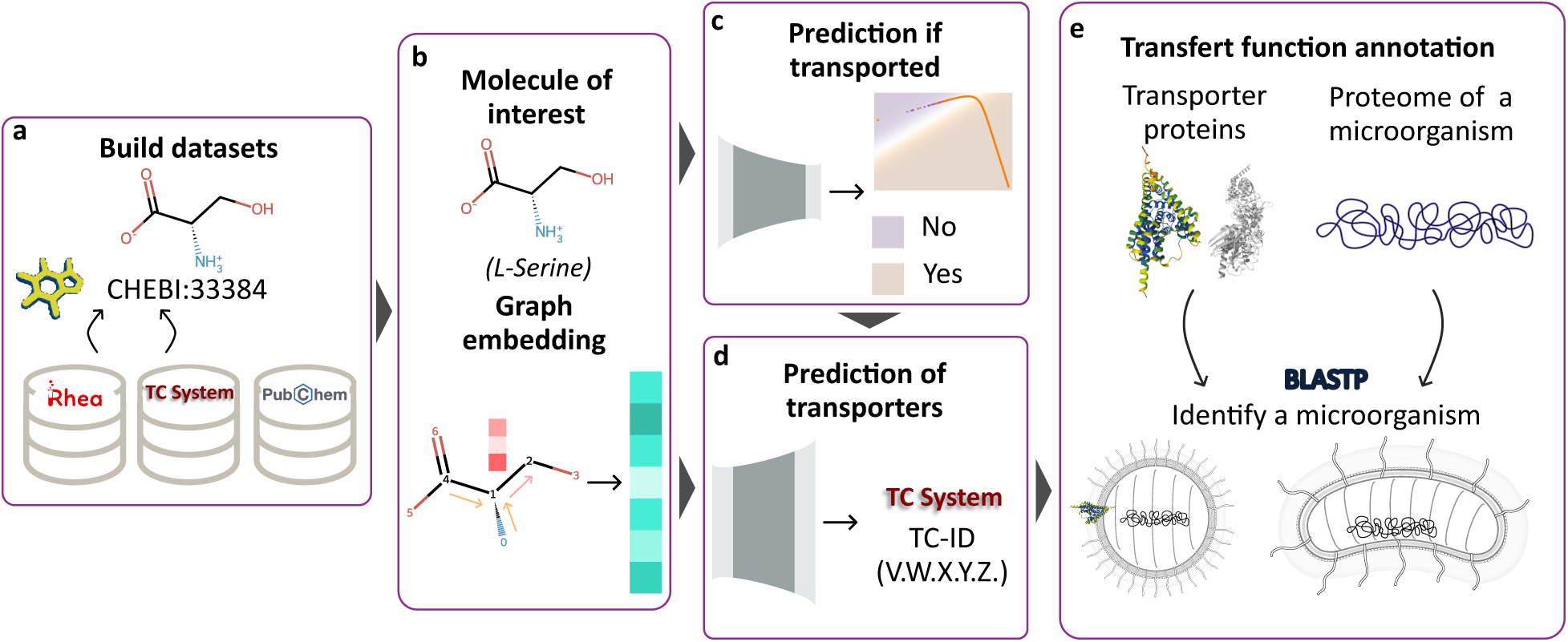
Model architecture. Once the datasets are constructed (**a**) the molecule of interest is encoded into a molecular embedding using the Directed Message-Passing Neural Network (D-MPNN) framework (**b**). This representation is then processed by the first module to determine whether the molecule is involved in a transport process (**c**). During the evaluation stage, if the molecule is predicted to participate in transport, the next module (**d**) infers its Transport Classification Number (TC-ID). Finally, based on the predicted TC-IDs, the corresponding protein sequences are used to identify microorganisms having transporters for molecular substrate (**e**).

### ChemProFlow predicts substrate involvement in transporter processes

Although existing databases catalogue molecules known to cross biological membranes, they provide no information about compounds that cannot undergo active transport. To address this limitation, we adopted a PU learning strategy, enabling the model to infer the ability of a molecule to traverse a membrane from partially supervised data. In this setting, known labeled samples are treated as positives, while remaining unlabeled samples may correspond to either class. PU learning is particularly well suited to such scenarios, where positive and negative examples are scarce. Among the commonly described PU learning approaches^25^, direct, two-steps, and PU bagging, the direct method was selected due to its conceptual simplicity and ease of integration within complex biological prediction pipelines (refer to “Positive-Unlabeled learning strategy” section of Methods). Accordingly, the classifier was trained using both positive and unlabeled samples, assuming a constant labeling probability across the positive class^30^. Here, positive molecules are those confirmed by two curated databases to cross membranes via carrier proteins, whereas unlabeled molecules are those for which transport capability remains unknown, allowing the model to learn discriminative patterns without requiring explicit negative annotations.

A TMAP visualization^31^ of molecular space (Figure 2a) highlights the distribution of labeled transport substrates versus unlabeled PubChem molecules. While unlabeled molecules form well-defined clusters, several appear in close proximity to characterized substrates. This spatial overlap suggests that some unlabeled molecules may indeed participate in transport processes but have not yet been experimentally validated, thereby supporting the suitability of PU learning for this task.

**Figure 2.**
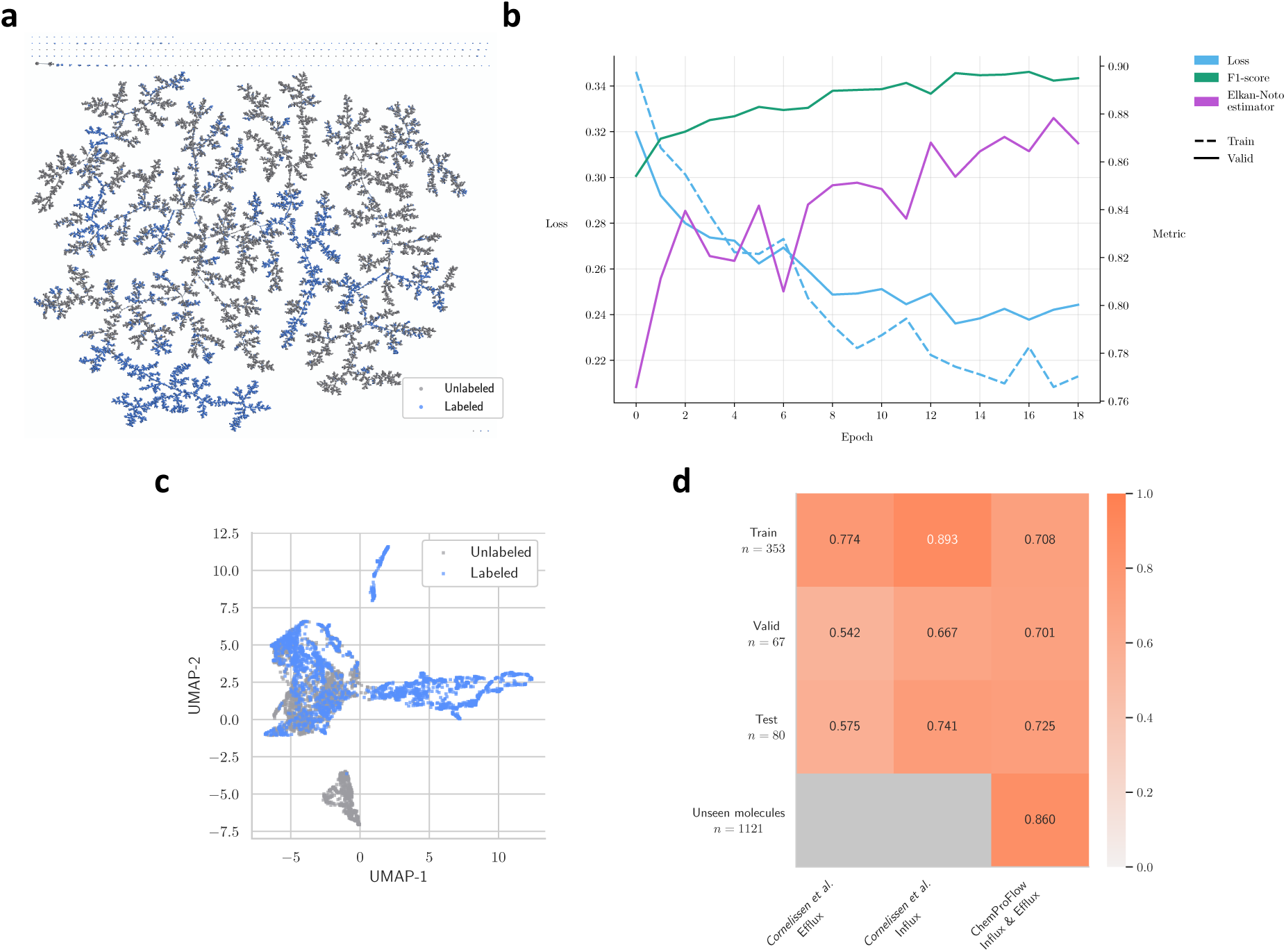
Representation learning and model calibration across labeled and unlabeled molecular data. **a** TMAP visualization of the molecular embedding space, illustrating the distribution of labeled and unlabeled samples used during training. **b** Training loss and validation performance metrics over epochs, showing model convergence and stability. **c** Two-dimensional projection of graph embeddings for 5,000 test molecules, revealing unlabeled instances located close to labeled examples and highlighting the overall structure and separation learned in the latent space. **d** Recall-based performance on the blood-brain barrier dataset, comparing ChemProFlow with an external framework and demonstrating the superior generalization capacity of ChemProFlow.

To discriminate between labeled and unlabeled molecules, we trained a GNN for which the binary classification performance is shown in Figure 2b (see “Training configurations” section of Methods). Training converged rapidly, with early stopping triggered at epoch 16 due to stable F1-score performance. Meanwhile, starting from epoch 12, the Elkan-Noto estimator, used to estimate the proportion of positives in the validation set, stabilized between 0.854 and 0.878 (mean = 0.867, standard deviation = 0.007), indicating convergence of the learning dynamics and consistency with the PU assumptions.

We next examined the graph embeddings learned by the model (Figure 2c). Embeddings corresponding to labeled molecules display substantial structural diversity and clear organization, whereas unlabeled molecules occupy a more compact region of the latent space. The partial overlap between the two groups reflects the intrinsic difficulty of distinguishing molecules with unknown transport properties and suggests that some unlabeled compounds may be plausible transport substrates.

To improve the reliability of predicted probabilities, we applied a post-hoc Dirichlet calibration model which adjusts the raw outputs of the classifier to better reflect true likelihoods^32^. Calibration substantially reduced the Binary Expected Calibration Error (Binary-ECE) from 3.92% to 0.37% and the Binary Maximum Calibration Error (Binary-MCE) from 16.82% to 7.99%, demonstrating markedly improved probability reliability (Supplementary Figure 1). Model performance on the test set also improved (Supplementary Table 1), yielding a more balanced trade-off between precision and recall and substantially limiting inclusion of molecules incorrectly predicted to cross membranes via transporter proteins despite lacking this capacity.

We further evaluated ChemProFlow on a challenging real-world benchmark: predicting molecular interactions with the Blood-Brain Barrier (BBB). BBB permeability is governed by several concurrent processes, including passive diffusion, endothelial transport, and transporter-mediated influx and efflux. We compared ChemProFlow against the tree-based model developed by Cornelissen *et al.*^33^, which models efflux and influx as independent tasks using molecular descriptors and fingerprints. In contrast, ChemProFlow handles both transporter types simultaneously through an inference mode that partitions outputs according to the requirements of each dataset. For molecules in the test dataset, ChemProFlow demonstrated a higher mean recall (0.725) than the tree-based model (0.658), as shown in Figure 2d. ChemProFlow showed superior generalization performance, particularly on molecules absent from the model development dataset, reaching a recall of 0.860 whereas the tree-based model exhibited a marked degradation in performance, with recall decreasing by 17.6% from the training to the test set. This performance underscores the capacity of the model to extrapolate beyond known substrates and capture the intricate molecular determinants governing transport across the BBB. However, because 27.70% of the molecules in our dataset are known to cross membranes yet are treated as negatives in the dataset of Cornelissen *et al.*, the false-negative rate cannot be reliably estimated in this comparison.

### ChemProFlow associates TC-IDs to a substrate

The second module introduces a novel approach that leverages molecular information to predict the TC-ID, analogous to enzyme classification efforts where EC numbers are inferred from biochemical reactions or protein sequences. A TC-ID comprises five hierarchical levels: the transporter class (TC-Class), subclass (TC-Subclass), family (TC-Family), subfamily (TC-Subfamily), and the specific transporter (TC-ID). As seen in enzyme classification studies, predictions are often made independently at each hierarchical level, as accuracy tends to vary between them. Within the ChemProFlow framework, one objective is to identify cellular transport mechanisms in microorganisms based on substrate characteristics. This raises the question of whether TC-ID prediction should be performed at a broad level, TC-class for example, or at the most specific transporter level. Because the fifth level of the TC-ID hierarchy corresponds to individual transporters, and therefore to specific microorganisms, we investigated whether truncating the TC-ID to the Subfamily or the class levels could still recover the same microbial associations. As shown in Figure 3a, when a TC-ID associated with a microorganism is shortened by removing lower levels of the identifier, the association retains up to 0.829 precision at the domain rank when using the TC-Subfamily level, and a precision of 0.002 at the species rank when using only the TC-Class level. These results suggest that only the complete TC-ID level preserves sufficient taxonomic information to reliably identify microorganisms possessing a given transporter mechanism, particularly at practical ranks such as species and genus.

**Figure 3.**
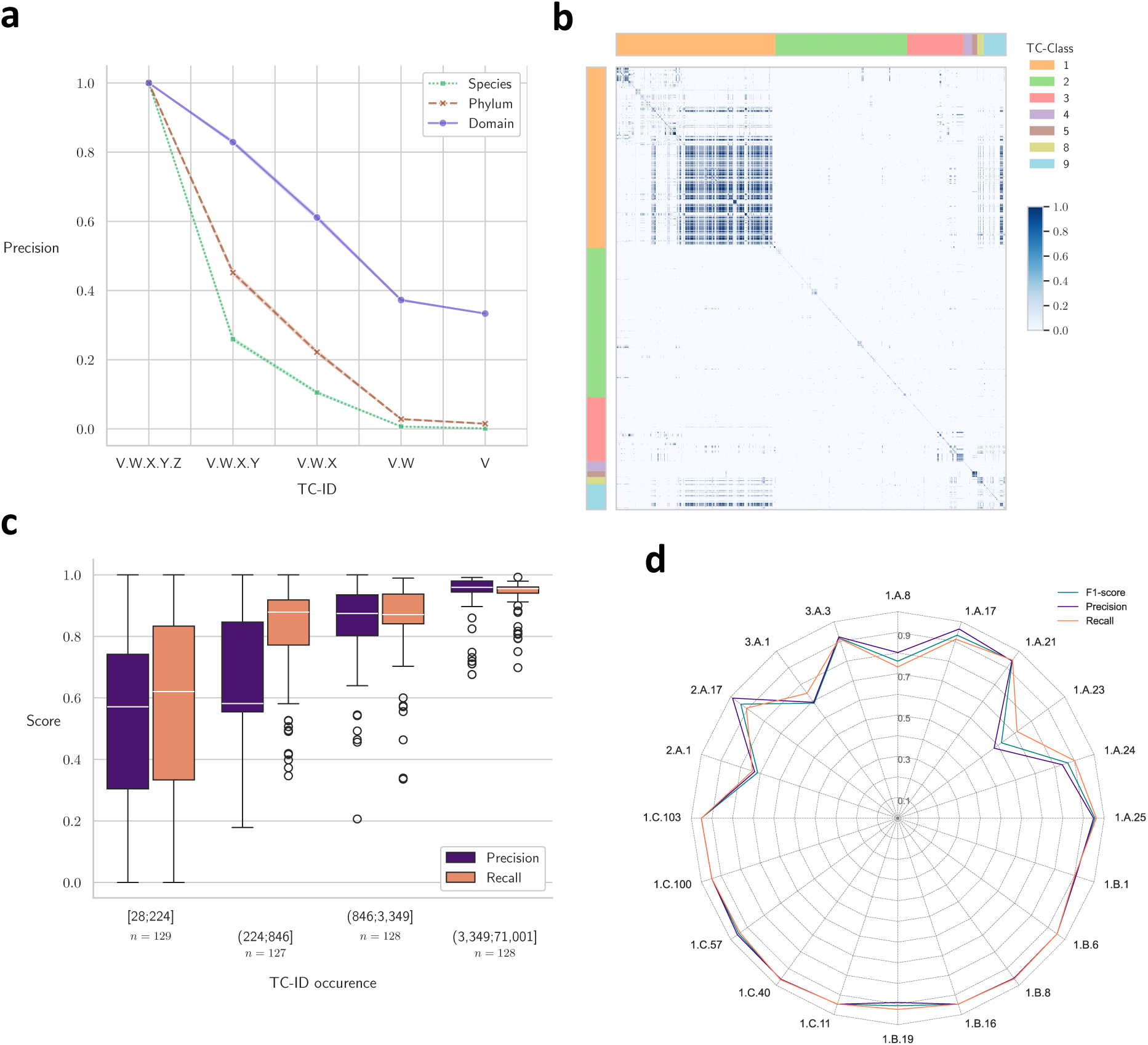
Evaluation of taxonomic retrieval, molecular similarity, and label distribution in transporter classification. **a** Precision of taxon retrieval across TC-IDs, illustrating accuracy at different hierarchical levels. **b** Hierarchical clustering of the Jaccard similarity between shared ChEBI compounds across TC-IDs, revealing functional and chemical relatedness among transporters. **c** Label classification performance stratified by label prevalence, showing how class imbalance influences predictive accuracy across categories. **d** Classification performance for the 20 most frequent transporter classes, illustrating model accuracy across the dominant TCID categories.

Different transporters, each identified by a unique TC-ID, can mediate the transport of the same substrate. Consequently, predicting TC-IDs from substrate representations constitutes a multi-label classification problem, since a single molecule may be associated with multiple transporters (see “Multi-label model” section of Methods). Although such problems can sometimes be reformulated as multi-class classification, preserving label associations during training provides a more accurate reflection of the biological relationships underlying transporter-substrate interactions^34^. Furthermore, several TC-IDs share identical sets of substrates, as illustrated in Figure 3b, where many TC-IDs from the first and ninth classes overlap substantially in their substrate profiles. To reduce learning complexity and avoid redundancy, we grouped TC-IDs that shared identical substrate sets and maintained a back-reference to the original identifiers. This strategy preserves the biological traceability of equivalent transporters while simplifying model training, reducing the total number of TC-IDs from 8,247 to 2,245 consolidated groups (Supplementary Figure 2).

As observed in functional predictions of enzymes or in biocatalytic predictions studies, some identifiers suffer from limited data availability and are typically excluded from model training and evaluation^9^. Following this principle, we removed substrates that were poorly represented and carefully addressed class imbalance among TC-IDs in the training and validation splits (refer to “Training configurations” section of Methods). As shown in Supplementary Figure 3, few labels remain underrepresented in the validation dataset, but all were retained in the test set to ensure fair evaluation.

Consistent with the substrate involvement prediction task, an early stopping mechanism was applied to prevent overfitting. The training and validation loss curves followed similar trajectories, while the F1-score stabilized around 0.925, achieving a final value of 0.927 (Supplementary Figure 4), in close agreement with the test dataset (Supplementary Table 1). Because of the inherent class imbalance, model performance was assessed using precision, recall, and F1-score, rather than relying solely on global accuracy^35^.

Beyond aggregate metrics, we also examined model performance on underrepresented labels. For TC-IDs represented by fewer than 225 molecules in the training set, precision and recall were 0.538 and 0.564, respectively, whereas for the most represented labels these metrics increased to 0.941 and 0.940 (Figure 3c). Across the 20 most frequent TC-classes, the model achieved a mean F1-score of 0.889 (Figure 3d). Although the underrepresented TC-IDs in the training dataset perform below state-of-the-art values reported for enzyme classification tasks^36^, the substrates associated with the more prevalent TC-IDs are consistently identified with high robustness.

We next assessed ChemProFlow in a challenging real-world context by introducing newly characterized substrates associated with defined transport mechanisms. As a case study, we focused on siderophores, molecules used in bioremediation^37^, and, more specifically, pyoverdines, which are produced by *Pseudomonas* species to scavenge iron from the environment. Although pyoverdine transport involves complex protein interactions^38^, the FpvA outer membrane transporter responsible for their uptake in *Pseudomonas aeruginosa* is recorded in TCDB (1.B.14.1.26), while its associated substrates are not explicitly annotated. Three characterized pyoverdine classes have been described. PvdI, PvdII, and PvdIII share conserved chemical motifs but differ in specific structural features that modulate their binding affinity to FpvA^39^. To assess the ability of ChemProFlow to generalize to unseen substrate variants within a shared transport mechanism, we incorporated PvdI and PvdII molecules (*n* = 55) into the training dataset and retrained the model. Notably, ChemProFlow correctly predicted all six PvdIII molecules, as susbtrates of the FpvA transporter (Supplementary Figure 5), demonstrating robust generalization across structurally related yet distinct siderophore classes. We then deployed the retrained model to screen the entire PubChem database (118,565,503 compounds) for candidates potentially transported via the FpvA mechanism. Altough none of the 61 curated pyoverdines were explicitly indexed as such in PubChem, ChemProFlow identified 1,275 compounds predicted to compatible with this transport mechanism. Among these, nine compounds carry the synonym “pyoverdine” or “pseudobactin”, supporting the biological relevance of the predictions and highlighting the capacity of ChemProFlow to recover mechanistically consistent candidates at scales.

### ChemProFlow searches protein transporters by homology

Homology-based searches for transporter proteins, such as those performed with G-Blast^12^ and gapseq^13^, typically rely on alignment coverage and scoring criteria derived from Blastp. These searches use an E-value threshold during alignment, combined with additional filters on alignment coverage and minimum sequence length. The RBH method is a well-established approach for identifying orthologous genes. In this context, transporter proteins can act either as query sequences, when genome proteins serve as the database, or as database entries, when genome proteins are used as the query. Previous studies on RBH searches have demonstrated the benefits of masking uninformative regions, applied Smith-Waterman alignment algorithms^40^, while others tested without composition-based score adjustment^41^. Moreover, the identification of the best hit can be based on either E-value or bit-score criteria^40^. Optimal parameter values depend on the software implementation, the size and composition of the database, and interactions among parameters. To identify the most suitable configuration for our application, we performed a systematic search across the parameter space identified from the literature (see “Reciprocal Best Hit parameters retrieval” section of Methods). Table 1 summarizes the parameters explored in our systematic optimization, their respective search ranges, and the worst- and best performing combinations obtained.

**Table 1.** Parameters and their search ranges used for Best Reciprocal Hit optimization, along with the final values selected after optimization. Each parameter is applied either during the Blastp search, the filtering of alignments results, or the selection of the best hit. The evaluation of the filtering locations as soft masks is active only if repeated sequences are filtered.

| Parameter | Context | Command-line argument | Values | Worst combination | Best combination |
| --- | --- | --- | --- | --- | --- |
| E-value | Blastp | -evalue | [10<sup>-12</sup>...10<sup>-3</sup>]<br>(step: x10) | 10<sup>-11</sup> | 10<sup>-5</sup> |
| Filter sequences with SEG | Blastp | -seg | Yes/No | No | No |
| Apply filtering locations as soft masks | Blastp | -soft_masking | Yes/No | No | No |
| Use local Smith-Waterman alignments | Blastp | -use_sw_tback | Yes/No | True | No |
| Use composition-based statistics | Blastp | -comp_based_stats | [0, 1, 2, 3] | 2 | 0 |
| Percent query coverage per hsp | Blastp | -qcov_hsp_perc | [20...90]<br>(step: 10) | 90 | 60 |
| Alignment length (%) | Filtering | / | [0...100]<br>(step: 10) | 60 | 50 |
| Selection of the best hit | Selection | / | Bit-score/E-value | E-value | Bit-score |

Across the 50 optimization iterations, only one parameter combination achieved the best F1-score of 0.788, while the average F1-score across all trials was 0.731. The lowest-performing configuration reached an F1-score of 0.569 highlighting the sensitivity of the RBH performance to parameter selection. This objective search strategy enabled identification of parameters that are highly data-dependent, particularly the E-value threshold. Notably, this optimization challenged previously reported key parameters in RBH applications, such as low-complexity sequences and the use of local alignment, which were not identified as critical for our specific application. Once the best set of parameters was identified, we compared the performance of the three approaches: our optimized RBH, GBlast, and gapseq, against the TC-IDs reported by Zafar and Saier^42^. In their original study, GBlast was used to identify transporter proteins within *Bifidobacterium* strains (see “Genome datasets” section of Methods). However, their workflow incorporated additional tools such as WHAT^43^ to evaluate protein hydropathy and HMMTOP^44^ to identify transmembrane regions, thereby refining the final transporter annotations through expert curation. A detailed comparison of the performance of the three methods across the nine *Bifidobacterium* strains is detailed in Table 2 (see “Comparison of homology-based search frameworks” section of Methods). Among them, GBlast achieved the highest mean retrieval rate of TC-IDs. However, gapseq produced a larger number of false positives, suggesting a tendency to over-predict transporter candidates when relying primarily on alignment-based heuristics and custom annotations, a behaviour that has already been reported^45^. In contrast, the optimized RBH approach demonstrated a strong balance between precision and recall achieving robust and reproducible performance without requiring manual filtering (Supplementary Figure 6). Moreover, as shown in Supplementary Figure 7, the distributions of the 10 most prevalent transporter classes are consistent between manual identification and RBH recovery across all nine *Bifidobacterium* strains. Overall, these results highlight that while expert-guided curation, can yield highly accurate transporter identification, automated and systematically optimized methods such as RBH can achieve good accuracy while maintaining scalability and reproducibility across multiple genomes.

**Table 2.** Mean performance metrics of gapseq, Gblast, and RBH across nine *Bifidobacterium* strains. Mean F1-score, sensitivity, and precision are reported for each method. Scores corrected for putative false positives are shown as primary values, with the corresponding raw scores provided in parentheses (see “Comparison of homology-based search frameworks” section of Methods).

|  | F1-score |  |  | Sensitivity |  |  | Precision |  |  |
| --- | --- | --- | --- | --- | --- | --- | --- | --- | --- |
|  | gapseq | Gblast | RBH | gapseq | GBlast | RBH | gapseq | GBlast | RBH |
| *B. adolescentis* ATCC 15703<br>(AP009256.1) | 0.315<br>(0.125) | 0.693<br>(0.655) | **0.810**<br>(0.779) | 0.333 | **0.974** | 0.887 | 0.298<br>(0.077) | 0.538<br>(0.494) | **0.746**<br>(0.695) |
| *B. animalis* subsp. *lactis* DSM<br>10140 (CP001606.1) | 0.275<br>(0.126) | 0.707<br>(0.676) | **0.835**<br>(0.804) | 0.301 | **0.962** | 0.885 | 0.253<br>(0.080) | 0.559<br>(0.521) | **0.790**<br>(0.736) |
| *B. bifidum* PRL2010<br>(NC_014638.1) | 0.278<br>(0.126) | 0.718<br>(0.674) | **0.800**<br>(0.780) | 0.280 | **0.959** | 0.808 | 0.276<br>(0.081) | 0.575<br>(0.520) | **0.792**<br>(0.754) |
| *B. breve* JCM 7017<br>(NZ_CP006712.1) | 0.329<br>(0.133) | 0.767<br>(0.717) | **0.839**<br>(0.807) | 0.295 | **0.970** | 0.859 | 0.371<br>(0.086) | 0.634<br>(0.569) | **0.820**<br>(0.761) |
| *B. catenulatum* DSM 16992<br>(NZ_AP012325.1) | 0.282<br>(0.125) | 0.737<br>(0.695) | **0.833**<br>(0.804) | 0.304 | **0.974** | 0.876 | 0.263<br>(0.079) | 0.592<br>(0.540) | **0.794**<br>(0.742) |
| *B. dentium* JCM 1195<br>(NZ_AP012326) | 0.341<br>(0.157) | 0.743<br>(0.713) | **0.838**<br>(0.821) | 0.333 | **0.952** | 0.864 | 0.350<br>(0.103) | 0.609<br>(0.570) | **0.814**<br>(0.781) |
| *B. longum* subsp. *infantis*<br>ATCC 15697 (NC_011593.1) | 0.337<br>(0.145) | 0.752<br>(0.704) | **0.819**<br>(0.788) | 0.315 | **0.969** | 0.825 | 0.362<br>(0.094) | 0.615<br>(0.553) | **0.812**<br>(0.754) |
| *B. longum* subsp. *longum*<br>JCM 1217 (AP010888.1) | 0.344<br>(0.149) | 0.745<br>(0.708) | **0.832**<br>(0.810) | 0.353 | **0.973** | 0.851 | 0.336<br>(0.094) | 0.604<br>(0.557) | **0.814**<br>(0.774) |
| *B. pseudocatenulatum*<br>(NZ_CP025199.1) | 0.312<br>(0.143) | 0.737<br>(0.710) | **0.846**<br>(0.827) | 0.314 | **0.955** | 0.865 | 0.311<br>(0.093) | 0.600<br>(0.565) | **0.828**<br>(0.791) |

### Assessing the performance of ChemProFlow

BioCyc is a comprehensive collection of Pathway/Genome databases that provides high-fidelity, manually curated annotations of metabolic and transport networks. Leveraging the Tier 1 and Tier 2 databases, we constructed a reference dataset of experimentally validated and highly reliable predicted transporters (see “Genome datasets” section of Methods). Analysis of this dataset revealed an average of 93.13 transported molecules per microorganism, yielding a total of 5,681 unique substrate-organism associations across the 61 bacterial species analysed. Among these, 753 molecules were unique, representing the distinct set of compounds predicted to participate in cellular transport processes (see “Genome scale metabolic model evaluation” section of Methods). This diversity highlights the chemical heterogeneity of transported substrates across microorganisms and underscores the broad functional coverage of the BioCyc-derived dataset. Using the RBH method, we identified a total of 4,667 unique TC-IDs across all microorganisms analysed. On average, each microorganism was associated with approximately 424 TC-IDs, representing the diversity of transporter systems detected within their proteomes. This extensive coverage reflects the high variability of transporter repertoires among species and supports the robustness of the RBH-based approach for large-scale identification of transport mechanisms.

For each microorganism, the ChemProFlow pipeline was executed on every molecule to determine whether it could cross the cellular membrane, to identify the corresponding transport mechanism, and to retrieve microorganisms predicted to encode that mechanism. Across the selected bacterial species, ChemProFlow correctly recovered the microorganism of origin for 89.53% (mean) of molecules not included in the curated dataset used for model development (Supplementary Figure 8). For molecules that were present in the model development dataset, the mean recovery rate was 82.83% (Supplementary Figure 9). In the two fungal species examined, the recovery rate reached 84.09% (Supplementary Figure 10), suggesting that although the RBH method has known limitations when applied to eukaryotic proteins with variable domain architectures^46^, it nevertheless maintains a high overall recovery in this context. These results demonstrate the strong ability of ChemProFlow to identify true transport-related instances across diverse microbial genomes. The consistently high sensitivity indicates that the integrated pipeline, combining molecular-level prediction, transporter classification, and ortholog detection, captures the majority of relevant transport mechanisms, underscoring the robustness and reliability of ChemProFlow for large-scale analyses of biological transport.

To illustrate the strengths and limitations of ChemProFlow, we analysed fluconazole, a commonly used antifungal drug. In the initial dataset, fluconazole was linked to eleven transporters across six microorganisms (Supplementary Table 2). After dataset curation, only four transporters remained, two of which, 2.A.1.2.123 and 3.A.1.205.32, were well represented, each associated with over 2,500 substrates. ChemProFlow correctly identified both transporters as candidates for fluconazole transport (Supplementary Table 3). The model also predicted additional transporter associations mediated through abstract ChEBI entities. For example, transporter 3.A.1.205.32 is annotated with azoles (CHEBI:68452), a parent chemical class that includes fluconazole and other related compounds that can serve as substrates. At the organism level, ChemProFlow identified *Candida albicans* and *Saccharomyces cerevisiae* as capable of transporting fluconazole (Figure 4a). This prediction was further supported by strong sequence similarity between the identified transporters and corresponding proteins in the BioCyc reference proteomes (see “Pairwise sequence alignment” section of Methods). While fluconazole was explicitly annotated only for *S. cerevisiae*, consistent with previous reports^47^, the prediction for *C. albicans* illustrates the ability of the model to infer plausible transport mechanisms beyond direct annotations, while also highlighting limitations arising from abstract substrate definitions and underrepresentation in curated datasets.

**Figure 4.**
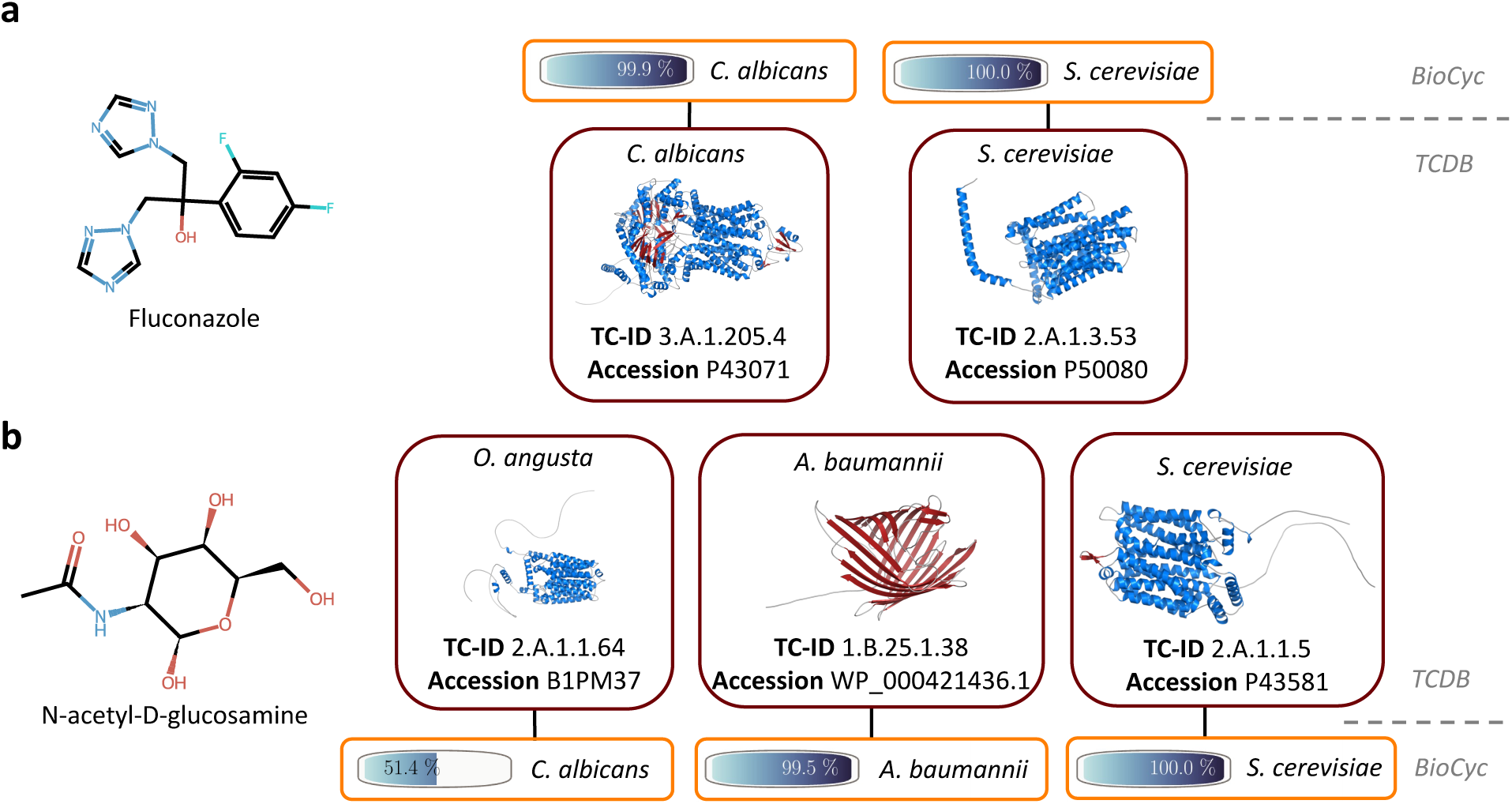
Identification of candidate microorganisms for substrate transport through predictive modelling of transport mechanisms and sequence homology. Microorganisms harbouring specific transport mechanisms are identified by first determining the transport mechanism and subsequently retrieving orthologous proteins from the microbial proteome using the Reciprocal Best Hit method. To assess result confidence, pairwise sequence similarity was calculated between the identified Transporter Classification Database (TCDB) entry and the corresponding BioCyc protein. **a** Two distinct mechanisms were identified for fluconazole transport, both exhibiting high sequence similarity between the TCDB reference and the BioCyc protein. **b** Of the three transport mechanisms identified for N-acetyl-D-glucosamine, two demonstrate high confidence matches between the TCDB and BioCyc databases, while the third indicates putative sequence homology between *Ogataea angusta* and *Candida albicans*.

To further assess the generalization capacity of ChemProFlow in a biotechnological context, we investigated N-acetyl-D-glucosamine, a value-added compound produced by engineered microbial cell factories. Membrane transporters play a central role in optimizing its production: deletion of uptake transporters enhances production yield, while the presence of dedicated export systems simplifies downstream processing^48^. This molecule, absent from the training dataset, is reported in BioCyc as transportable in *Escherichia coli* B str. REL606. ChemProFlow correctly identified N-acetyl-D-glucosamine as a candidate for transport but did not recover this specific strain, despite prior experimental evidence of its transport capability^49^. Nevertheless, the model identified several plausible transporter mechanisms (Supplementary Table 4). Notably, as shown in Figure 4b, ChemProFlow predicted transport via the membrane porin OprD (1.B.25.1.38) from *Acinetobacter baumannii*. OprD is homologous to the porin characterized in *P. aeruginosa*^50^ which has been experimentally shown to transport N-acetyl-D-glucosamine^51^. The model also identified transport through a native hexose transporter (2.A.1.1.5) in *S. cerevisiae*, consistent with published uptake studies^52^. Furthermore, ChemProFlow identified the equivalent transporter 2.A.1.1.64, which shares an identical substrate profile and is present in *Ogataea angusta*. This transporter shares 54.1% sequence similarity with the glucose transporter of *C. albicans*. The homology between this transporter and the *S. cerevisiae* hexose transporter provides evidence supporting the functional relevance of the identified mechanism^53^.

Together, these results suggest that ChemProFlow can extrapolate beyond curated annotations to recover biologically plausible transporter-substrate-organism associations. On average, the pipeline recovered 6.63-fold more candidate transportable molecules per microorganism than those reported in the original BioCyc dataset (Supplementary Table 5). While these additional predictions do not necessarily imply confirmed transport activity, they highlight the capacity of the framework to uncover previously unannotated but biologically plausible transport relationships, providing a basis for targeted experimental validation.

## Discussion

Understanding how molecules are transported across biological membranes remains a fundamental challenge in biology. While decades of research have catalogued transporter proteins, the field has largely remained protein-centric, inferring function from sequence similarity. In contrast, the molecular determinants that govern whether a compound can be actively transported and by which mechanisms have remained comparatively underexplored. Here, we introduce a complementary, substrate-centric framework that shifts the perspective from “which proteins transport?” to “which molecules can be transported, how, and by whom?”. By leveraging the hierarchical structure of the Transport Classification Database (TCDB) and integrating Geometric Deep Learning (GDL) with positive-unlabeled learning and orthology-based mapping, ChemProFlow establishes a direct link between chemical structure, transport mechanism, and genomic context. This unified framework enables three levels of inference: predicting whether a molecule is actively transported, identifying the most likely transporter class, and mapping these transport capabilities across microorganisms.

Many of the challenges faced when inferring enzyme function and predicting biochemical reactions using EC numbers are also encountered when TC-IDs are used as intermediaries to describe transporter activity. Recent advances have focused on improving molecular representations and classification strategies to overcome the intrinsic limitations of such datasets. For instance, GDL approaches have been enhanced with molecular fingerprints, integrating domain-specific chemical knowledge into molecular embeddings to improve predictive accuracy and interpretability^54^. To address data imbalance, contrastive learning frameworks have been shown to sustain high performance for underrepresented biochemical reactions^55^, while additional specialized techniques further enhance classification robustness^56,57^. Future work will leverage the modular architecture of ChemProFlow to incorporate such advancements into each component independently.

Prediction granularity across the TC-ID hierarchy presents an additional trade-off. While family-level, third-level, assignments often provide greater robustness^58^, complete TC-ID resolution is required to compare the presence or absence of specific transport mechanisms across microorganisms. As transporter databases catalog transport proteins across diverse taxa, their continued expansion and refinement will be essential for improving high-resolution transport mapping^59^. Achieving such high-resolution comparisons across microorganisms depends on accurate ortholog detection within genomic datasets. The Reciprocal Best Hit (RBH) method, although widely used for ortholog inference, has known limitations, including restricted scalability, difficulty handling many-to-many relationships, and limited capacity to account for gene losses or complex genomic rearrangements that may generate artefactual matches^15^. In our context, these limitations are partially mitigated by the focus on predominantly one-to-one transporter relationships. Nevertheless, integrating more sophisticated ortholog detection approaches could further reduce false-positive predictions and improve transporter identification precision. Objective optimization of RBH parameters further revealed that performance is highly data-dependent. Interestingly, parameters frequently emphasized in prior RBH applications, such as low-complexity sequence filtering or the use of local alignment, did not substantially influence performance in our specific setting. These findings suggest that optimal RBH parameterization may vary depending on dataset composition and biological context, underscoring the importance of context-aware optimization rather than reliance on standard heuristics. Importantly, the presence of a transporter gene in a microorganism does not necessarily imply its systematic expression or functional activity under all conditions^2^. Our strategy identifies genomic transport potential, which may not fully reflect context-dependent transcriptional or post-transcriptional regulation.

Beyond methodological advances, the substrate-centric paradigm introduced here has broader implications. In pharmacology, it may facilitate early-stage assessment of compound transportability and transport mechanisms. In bioremediation, mapping transport capabilities across taxa offers a new lens for investing the diversity of substrate uptake across microbial communities. In bioproduction, identifying the transport mechanisms associated with a target molecule can guide strain engineering strategies, such as modulating uptake or export systems, to enhance product yield. More broadly, integrating chemical structure, transport mechanism, and genomic context contributes toward a systems-level understanding of membrane transport as a dynamic interface between organisms and their chemical environment.

In summary, ChemProFlow provides a modular and extensible open-source framework that bridges chemical space and genomic transport capacity. By unifying graph-based molecular learning with orthology-based mapping, it establishes a scalable foundation for investigating membrane transport from the substrate perspective. We anticipate that this conceptual and computational integration will enable new research directions across pharmacology, biotechnology, and microbiology.

## Methods

### Transport substrate datasets

We processed the Rhea^60^ database (release 139) from the Biological Pathway Exchange file to extract the molecules, referenced by their ChEBI^61^ identifiers, involved in a transporter processes, along with notes about their cellular locations (Supplementary Note 1). After associating the Uniprot^62^ identifiers with these Rhea reactions, we linked the TC-IDs to these sequences using the TCDB database. We retained only the substrates with an associated TC-ID, resulting in 4,878 TC-IDs linked to ChEBI entries. We then merged this set with the *get_substrates.tsv* file provided by TCDB which contains 14,317 TC-ID-ChEBI associations, to create a dataset of 19,051 tuples (Supplementary Figure 11).

Since a ChEBI identifier can refer to abstract molecules, we expanded each ChEBI entry (*n* = 1,943), into its corresponding real molecules using the ChEBI ontology. However, some molecules contain R-groups, which act as placeholders for variable molecular fragments. To replace these undefined components, we substituted each R-group structure with existing molecules sharing the same core scaffold, using the BioRGroup^63^ dataset as the source of template-based expansions. To ensure that the stereochemistry was fully defined in the molecular representations, we selected the first stereoisomer of each molecule generated by the *EnumerateStereoisomers* module in RDKit^64^ (v2024.9.3). In addition, formal atomic charges were retained, as they provide chemically relevant information used later by the model. The final dataset establishes 162,198,714 associations between transported substrates and their corresponding TC-IDs, encompassing 155,251 unique molecules and 9,082 unique transporters.

Finally, to construct the unlabeled dataset, we retrieved the full PubChem^65^ FTP archive (accessed on 2024/08/03) and integrated it into an in-house chemical database. We then selected molecules with a natural-product likeness score greater than zero, since structures enriched in natural-product features are more likely to be metabolically relevant, and thus more susceptible to be encountered and transported within living cells (Supplementary Figure 12). This procedure yielded a dataset comprising 151,238 transported substrates and an equal number of unlabeled molecules, enriched for natural-product-like compounds that are difficult to distinguish from true transport substrates. For the second dataset used to predict TC-IDs from molecules, we retained only TC-IDs associated with at least 50 molecules, and we removed duplicated groups of substrates associated to each TC-ID. Because this deduplication step is inherently arbitrary, we preserved a mapping of merged TC-IDs to maintain traceability. The resulting dataset contained 512 unique TC-IDs associated with 151,114 unique molecules, with a mean label cardinality of 22.66. Pyoverdines from three distinct classes, as described by Cezard *et al.*^66^ were included in this study (Supplementary Table 6). Of the 66 molecules originally reported, 61 were retained for analysis (classe I, *n* = 20; class II, *n* = 35; class III, *n* = 6), corresponding to compounds for which both the pyoverdine sequence and chemical structure were consistently matched.

### Genome datasets

A collection of nine *Bifidobacterium* strains was obtained from Zafar and Saier^42^. The dataset includes the TC-IDs associated with each strain, and their corresponding proteomes were retrieved from the NCBI web service^67^. The TC-IDs from *B. longum* subsp. *infantis* ATCC 15697 were used to optimize the parameter combinations for the Reciprocal Best Hit (RBH) procedure.

GEMs that had at least one year of supporting literature or were manually curated were downloaded from BioCyc^68^ (v29.0). The reconstruction of the full lineages was achieved through the utilisation of the Taxonomy database^69^, which had been integrated into a built-in house database in advance. Bacterial and fungal microorganisms were selected based on the availability of a proteome and their taxonomy identifiers: 2 at the domain level for Bacteria and 4751 at the kingdom level for Fungi, resulting in 61 bacterial and 2 fungal models.

### Molecular embedding

Directed Message-Passing Neural Networks (D-MPNN) have proven effective for molecular property prediction. A D-MPNN operates in two main phases: the message-passing phase, which constructs molecular latent representations, and the readout phase, which produces task-specific predictions. The message-passing mechanism is described in this section, while the readout phase is detailed in the corresponding model sections.

Each atom and each bond in a molecule is represented as a feature vector following the same encoding scheme used in Chemprop^29^. During message passing, edge-level hidden states are iteratively updated through an edge-centric propagation scheme composed of three message-passing layers, enabling the model to integrate information from progressively larger local neighbourhoods. At each layer, messages from adjacent directed edges are aggregated using a mean operator. After the message-passing phase, atom-level hidden states are obtained by summing the incoming edge representations for each atom. In the readout stage, these atom-level hidden states are further aggregated using a sum operation to produce the final molecular embedding used for downstream prediction tasks. This core model was implemented using the PyTorch Geometric (v2.7.0)^70^ library.

### Training configurations

Models were trained using a K-Fold cross-validation strategy, which splits the dataset into 𝒦 folds (𝒦 = 5). To ensure balanced distributions of classes or labels, we used either the StratifiedKFold^71^ or MultilabelStratifiedKFold^72^ approach, depending on the task. All models were implemented with PyTorch (v2.9.1) and Lightning (v2.5.5)^73^, which provides standardized machine learning training routines. Performance metrics, including precision, recall, and F1-score, were monitored using TorchMetrics^74^.

The models were trained with an initial learning rate value of 10^−3^ using the Adam optimizer; and a batch size of 16. To prevent overfitting, early stopping was applied: training stopped when the validation F1-score failed to improve by more than 5e^−3^ over five epochs. Among the five models produced by the five-fold training procedure, the best-performing model on the test set was retained.

### Positive-Unlabeled learning strategy

The readout component of the model consists of standard Feed Forward Network (FFN) with ReLU activation, mapping the atom-level hidden states to a single output neuron. The model is trained using a Binary Cross-Entropy (BCE) loss, which measures the discrepancy between predicted and expected class probabilities, and this loss is used to backpropagate the gradients.

To implement the PU learning workflow, we made several assumptions about the dataset. The dataset consists of molecules involved in transport processes, which are considered positive data points, molecules from PubChem that resemble natural products, which are considered unlabeled data points. First, we assumed that both the created dataset has positive and unlabeled data coming from the same dataset. This assumption is reasonable because the unlabeled data were selected based on their natural-product likeness, ensuring chemical similarity with the molecules found in a microorganism and carry-out by a transport mechanism.

Second, we assumed that the set labeled molecules represent a uniform subset of all positive examples, meaning that the labeled samples were Selected Completely At Random. Under this assumption, the propensity score:

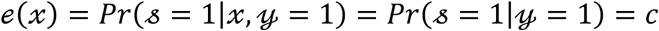

is constant and corresponds to the label frequency *c*. Here, 𝓈 = 1 denotes a labeled sample, 𝓎 = 1 denotes a positive sample, and 𝓍 represents the molecular features. Consequently, the probability of a sample being labeled is proportional of it being positive, scaled by the label frequency *c*^25^:

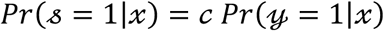

The label frequency *c* was estimated as the average predicted probability across the labeled validation dataset^30^. A Dirichlet post-hoc calibration was then applied using a logistic regression model trained on the predicted probabilities *Pr*(𝓎|𝓍). The optimal classification threshold was selected from the validation set by scanning values from 0.01 to 0.99 (increments if 0.01) and choosing the cutoff that maximized the F1-score.

To evaluate calibration quality, we computed two metrics: the Binary-ECE, which measures the mean discrepancy across bins in the reliability diagram, and the Binary-MCE which measures the largest such discrepancy^75^. These metrics are defined as follows:

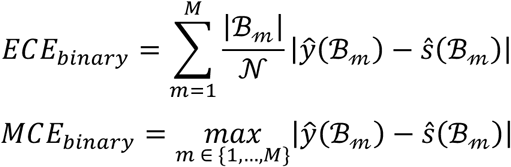

Where ℳ denotes the number of bins, ℬ_𝓂_ the bins under consideration, |ℬ_𝓂_| the number of bins, ŷ(ℬ_𝓂_) the proportion of positives, *ŝ*(ℬ_𝓂_) the average probability, and 𝒩 the number of instances.

### Multi-label model

The readout component of the model is implemented as a FFN with ReLU activation, which maps the atom-level hidden states to an output layer containing 512 neurons, corresponding to the number of labels. The model is trained using the BCE loss function, which quantifies the discrepancy between the predicted and true label probabilities for each output.

### Reciprocal Best Hit parameters retrieval

To identify orthologs as RBHs, we verified, for each *best hit* found by a query protein in the forward search, whether the corresponding target gene also identified the query as its best hit in the reverse direction^76^. Although Blastp is not the fastest alignment tool available, it remains a benchmark method for this type of analysis^77^. However, its configuration can significantly influence the results and may lead to unexpected outcomes if not carefully tuned^78^.

The RBH procedure involves three main stages: (i) configuring the Blastp parameters, (ii) filtering to retain only the best hits, and (iii) selecting reciprocal matches. To determine the optimal combination of settings and parameters values, we performed a systematic optimization using Optuna^79^, a library for automated hyperparameter tuning, based on the Tree-structured Parzen Estimator algorithm. In this context, the objective function to be optimized corresponded to the RBH detection process, evaluated using the TC-IDs retrieved from the proteins associated with each best hit compared to those identified by Zafar and Saier^42^. The F1-score served as the optimization criterion and was maximized over 50 iterations, during which Optuna explored different parameter combinations to identify the best-performing configuration.

### Comparison of homology-based search frameworks

We compared Gblast (https://github.com/SaierLaboratory/TCDBtools, accessed August 2025), gapseq (v1.4.0), and our optimized RBH method across the nine *Bifidobacterium* strains. For each method, we quantified the number of TC-IDs retrieved and compared them with the reference transporter annotations. For gapseq, the original manually curated transporter protein lists and annotation files provided by the authors were used for evaluation. In contrast, for Gblast and the RBH approach, transporter proteins identified in this study were used as input. All software tools were executed using their default parameter settings, except where explicitly noted for the optimized RBH method.

In a secondary evaluation step, candidate transporters identified in the genomes but absent from the original study were not automatically considered false positives if they carried functional annotations indicative of transport activity. Specifically, proteins annotated with terms such as “aquaporin”, “antiporter”, “symporter”, “transporter”, “channel”, “permease”, “efflux pump”, “ABC exporter”, “solute carrier”, “exporter”, or “transporter system” were retained as plausible transporter candidates.

### Pairwise sequence alignment

Global sequence alignments were performed using the Needle program via the EBI web service^80^ with default parameters (accessed January 2026).

### Genome scale metabolic model evaluation

In the first step, we selected the molecules involved in transport processes for each microorganism from the BioCyc database. We parsed the associated ontology to identify reaction identifiers annotated as *Transport-Reactions* in their *type* fields or containing *IUBMB* references in their *comment* sections, while explicitly excluding reactions labeled as *Simple-Diffusion* in their *common name* fields. Reactions were retained only if their associated metabolites were annotated with compartment information, specifically *CCO-IN* or the *CCO-OUT*, indicating the intracellular or extracellular location of the molecule, respectively. Molecules were then selected if they appeared in both compartments and participated exclusively on one side of the reaction, suggesting a directional transport role. Each selected molecule was mapped to its corresponding compound identifier using the BioCyc compound file and retained only if its associated SMILES string could be successfully parsed using RDKit, ensuring structural validity for downstream modelling.

In the second step, we identified the transporter proteins for each proteome using the RBH method with the optimized parameter set determined previously. Finally, in the third step, we applied the full ChemProFlow pipeline to each molecule: first determining whether the molecule was predicted to participate in a transport mechanism, if so, retrieving the associated TC-IDs, and subsequently identifying the microorganisms containing the corresponding transporter mechanisms.

### Dataset characterization and model evaluation metrics

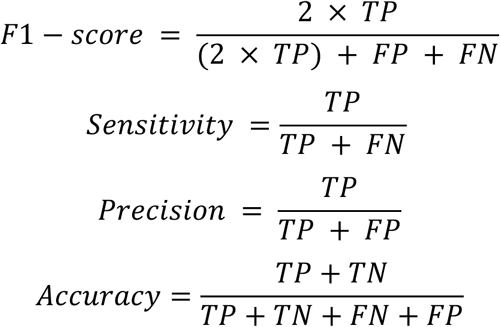

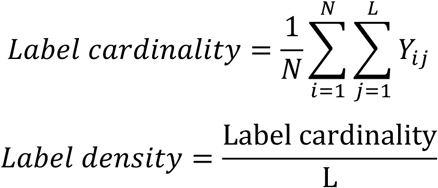

With:

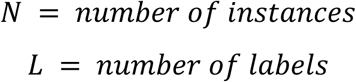

## Data availability

Datasets and trained models are available at Recherche Data Gouv (https://doi.org/10.57745/QXBLVM).

## Code availability

The code used for dataset construction, as well as for model training and inference, is available on GitHub (https://github.com/brsynth/chemproflow) and Zenodo (10.5281/zenodo.18913584).

## Acknowledgments

This work was supported by a French government grant managed by the Agence Nationale de la Recherche under the France 2030 program, reference ANR-22-PEBB-0008. We are grateful to the Genotoul bioinformatics platform Toulouse Occitanie (Bioinfo Genotoul, https://doi.org/10.15454/1.5572369328961167E12) for providing computing and storage resources. This project was provided with computing HPC and storage resources by GENCI at IDRIS thanks to the grant AD010314674 on the supercomputer Jean Zay’s V100 partition. Part of the computing analyses were also performed on the Core Cluster of the Institut Français de Bioinformatique (IFB) (ANR-11-INBS-0013).

## Author contributions

G.G. (Conceptualization, Data Curation, Investigation, Methodology, Software, Validation, Visualization, Writing - original draft), T.D. (Conceptualization, Funding acquisition, Resources, Writing - review & editing), P.M. (Conceptualization, Writing - review & editing), and J-L.F. (Conceptualization, Funding acquisition, Project administration, Writing - review & editing).

